# Is the blunderbuss a misleading visual metaphor for stasis and punctuated evolution?

**DOI:** 10.1101/242297

**Authors:** John T. Waller

**Affiliations:** Evolutionary Ecology Unit, Department of Biology, Lund University, SE-223 62 Lund, Sweden

## Abstract

I discuss the usefulness of the so-called “blunderbuss pattern” of phenotypic evolution as a visual metaphor for stasis and punctuated evolution that was originally put forward in Uyeda et al. (2011) in their highly influential paper “The million-year wait for macroevolutionary bursts”. I argue the blunderbuss pattern is not surprising, and in some cases it is misleading. I review several publications that cite Uyeda et al. (2011) that seem to be confused about the meaning of the pattern and what it implies. I do not critique the original analysis within Uyeda et al (2011), but show the blunderbuss pattern itself would be produced even when assuming a Brownian motion (completely gradual) model of phenotypic divergence. Finally, I discuss how the interesting results of the paper have been overlooked in favor of the surprisingly powerful, but also misleading visual metaphor of the blunderbuss.

## Introduction

In a highly influential and impressive macroevolutionary study, Uyeda et al. (2011) presented compelling evidence that body size evolution occurs in rare but substantial bursts, with bounded evolution on shorter time scales (Uyeda et al. 2011; their Table 1). These authors fitted several models to various sources of body size divergence data, compiled from fossil evidence, phylogenetic comparative methods, and studies of contemporary evolutionary rates. They found that the best-performing model was the multi-burst model of evolution. Furthermore, the parameter estimates from this model indicated that there is at least a 1 MYA wait for these bursts of body-size evolution. These significant and exciting results were presented in conjunction with a powerful visual metaphor that has also became highly influential: “the blunderbuss” (Uyeda et al. 2011, Fig. 1).

**Figure 1.**
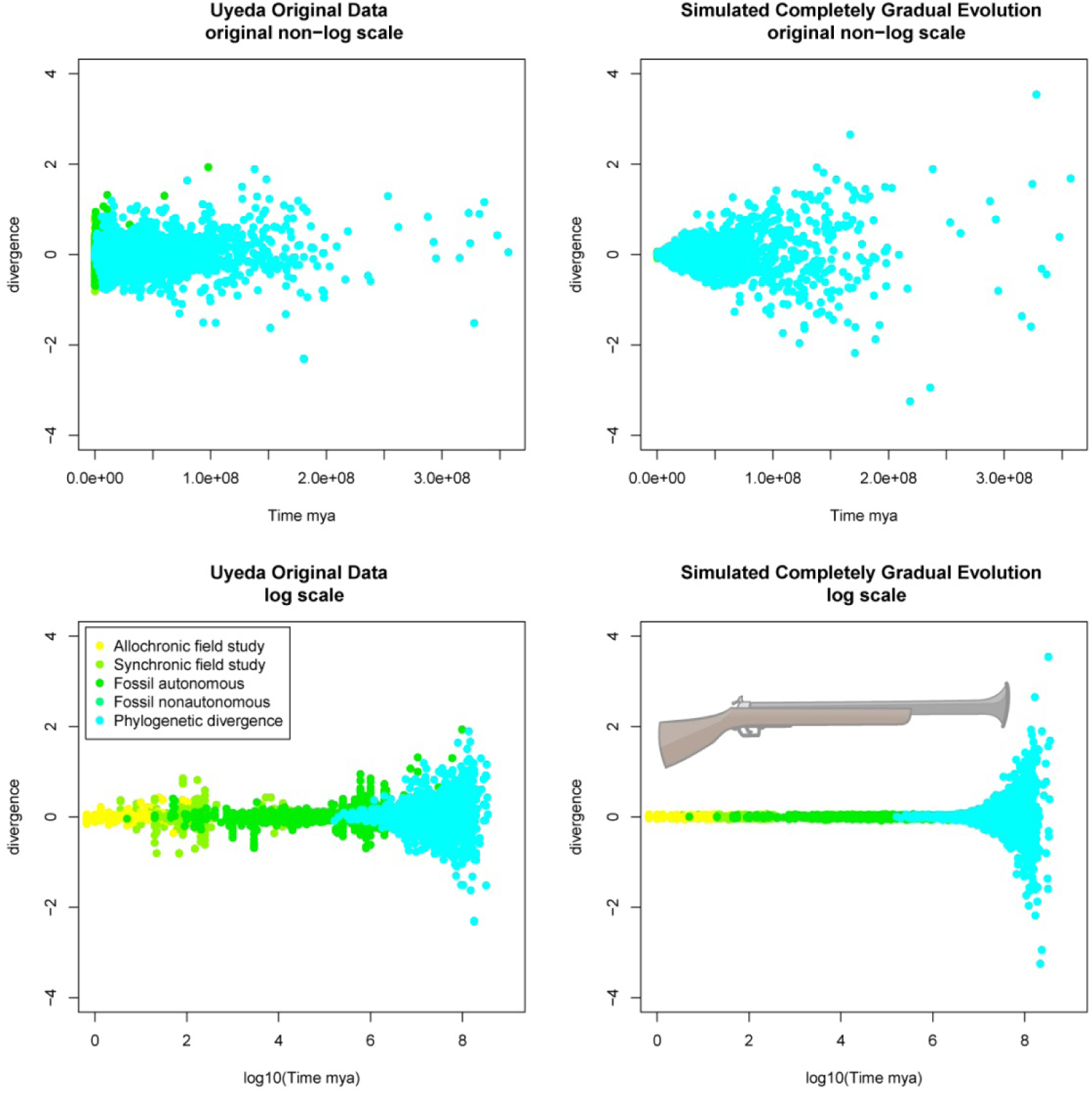
Upper left panel, original data from Uyeda et al. (2011) plotted on original non-log scale. Here we see that much of the divergence data is not interpretable because divergence time is less than 100,000 years, explaining why Uyeda et al. would choose to plot it on a log scale in the lower left panel. In the upper right panel, I have used the time of divergence from the original Uyeda et al. (2011), but instead of using the empirical divergence, I have simulated divergence using a completely gradual model of evolution (Brownian motion). In the lower right panel, I have plotted this simulated gradual evolution on a log scale and a very distinctive blunderbuss pattern appears. From this plot it is clear that the logging of the y-axis causes all the data to be squished into the final third of the plot, resulting the flared blunderbuss pattern even when phenotypic evolution is entirely gradual. An actual blunderbuss is also shown as a reference.Potentially, the blunderbuss illustrates the “long barrel of stasis,” but we see also that the BM-model can also produces a similar long barrel of stasis (Fig. 1; lower right panel) even though under this model, evolution occurs gradually and variance in phenotypic space is accumulating at a linear rate. Any appearance of bounded stasis in a Brownian motion model is therefore entirely due to it being plotted on a log scale. So if the visual metaphor of the blunderbuss as stasis cannot easily be distinguished from Brownian motion, is this metaphor even useful or is it simply misleading?

Uyeda et al. (2011) also tested an appropriate null model, Brownian Motion (BM), which makes very different predictions from their best performing model (the multi-burst). Under the BM-model, evolution is unbounded and occurs gradually (i.e. no paradoxical 1 MYA waits for bursts of evolution). Instead, under such a BM-model, phenotypic evolution accumulates continuously and gradually (Felsenstein 1985; Hansen 1997). These two models tell very different stories about the how body size diverges over time.

Here, I present the results from a simulation of divergence under a BM-model (Fig. 1). Interestingly, and somewhat problematically, a BM-model will also produce a blunderbuss pattern when plotted on a log scale. Thus, even though the BM-model and the multi-burst model (constrained evolution below a million-year bound) are qualitatively and quantitatively very different models of evolution, both types of models will produce a blunderbuss pattern on a log scale. One might therefore legitimately worry and wonder what is so special or instructive about a blunderbuss pattern? In other words, no fundamentally novel insights into any of these two models of evolution can be gained from the visual discovery a blunderbuss pattern in your data. Thus, the fact that the visual pattern that is observed happens to be resemble a blunderbuss when plotted on a log-scale is neither special, surprising, or provides any interesting insight.

This commentary is not meant to be a criticism of the original paper of Uyeda et al (2011) nor of its methods. The intention of this commentary is only to draw attention to the fact that the blunderbuss pattern of body size divergences that was plotted in Uyeda et al. (2011, Fig. 1) is neither surprising or unique to their model or has any real biologically meaningful interpretive value. I therefore merely want to point out that the so-called blunderbuss pattern is almost guaranteed result when a class of closely related models of phenotypic evolution plotted on a log scale. The non-exhaustive list of interesting aspects of Uyeda et. al. (2011) are as follows:

1. The large dataset used.
2. The large temporal scale.
3. The diversity of taxonomic groups used.
4. The fact that the best-fitting model was a multi-burst model.
5. That the model parameters were consistent across several taxonomic groups.

The non-interesting aspects of Uyeda et al. (2011) is the fact that phenotypic divergence looks like an old-fashioned gun pattern when plotted on a log scale. However, this visual pattern seems to be the aspect that has captured the attention of other evolutionary biologists, and that has consistently and often been emphasized in many discussions and citations of the paper. For example, Voje (2016) cites the blunderbuss pattern as evidence that lineages evolve in narrow zones:

> “That most traits show little net evolution but still undergo substantial change is consistent with the hypothesis that lineages mostly evolve within rather narrow adaptive zones, as envisioned by Simpson (1944, 1953) and as observed by Uyeda and colleges in the blunderbuss pattern of body size evolution.” p. 2685 par. 2

Doebeli and Ispolatov (2016) go even further and refer to a “blunderbuss theory” of temporal patterns of macroevolutionary changes and diversification, but later more correctly characterize the results of the multi-burst model (Uyeda et al. 2011). But is the multi-burst model really best characterized as the “blunderbuss theory” when Brownian motion also produces a blunderbuss pattern? Fortelius et al. (2014) see the “the blunderbuss pattern” as compatible with the punctuated equilibria model of (Eldredge and Gould 1972), but never really discuss the multi-burst model.

Arnold (2014), who was one of the co-authors of Uyeda et al. (2011), recognizes that simple a simple BM-model can produce a blunderbuss pattern, but falls short of rejecting it as a useful visual metaphor as I do here (p. 738 para. 5). Later, Arnold (2014) also recognized that barrel of the blunderbuss does not represent absolute stasis:

> “The bounded evolution revealed by the slender barrel of the blunderbuss should not be confused with stasis. The limits of that slender barrel lie 6 within-population standard deviations, or a 65% change in body size, on either side of literal stasis. Within those bounds, individual species can appreciably evolve.” p. 742 para. 5

Finally, even the first author of Uyeda et al. (2011) seems to be aware that the blunderbuss might have been taken the wrong way. In the comment section of Jerry Coyne’s blog “Why Evolution is True” (https://whyevolutionistrue.wordpress.com/2011/09/22/want-evolutionary-change-wait-a-million-years/) about the article “Want evolutionary change? Wait a million years” user Uyeda writes:

> “The blunderbuss pattern itself isn’t the product of punctuated equilibrium certainly. It is what you would see in either the case of a gradual Brownian motion or repeated punctuations (our Multiple-Burst model) on the log timescale (see Figure 3 in our paper) Consequently, it’s not surprising that it eventually looks like a blunderbuss, what was impressive to us was the consistency of pattern, timing and parameter estimates between independent data sources across a wide range of taxa.” (https://whyevolutionistrue.wordpress.com/2011/09/22/want-evolutionary-change-wait-a-million-years/#comment-135938)

How useful is the blunderbuss as a visual metaphor for stasis or punctuated evolution? Here, I have argued that it varies from being misleading to technically correct, but for the wrong reasons. Often a creative use of new terms and metaphors can be a good way to get a concepts to stick in the minds of the reader, but I think in this case the mental imagery has been too strong. The metaphor of the blunderbuss pattern has caused many readers to believe it is mainly the visual pattern that is interesting, thereby drawing attention away from the other results of the paper. The fact that completely gradual evolution can produce a blunderbuss pattern, casts doubt on the metaphor as a useful mascot for stasis and punctuated evolution.

## Methods

All analysis was done in R. R-code can be found in the supplementary material. I downloaded the dryad package of Uyeda et al. (2011) (doi:10.5061/dryad.7d580). Using both the “Dryad7.csv” and “Phylogeniesbynode.csv”, I extracted divergence times and phenotypic divergence data. Using this data, I re-plotted the original Figure 1 in Uyeda et al. (2011). I did not, however, include the phylogenetic time-series data, so the two figures will appear slightly different. Using the divergence times from the original study, I simulated simple phenotypic evolution by Brownian motion in R (σ = 0.001). I simulated 5000 independent runs of phenotypic divergence under a Brownian motion model. In each model a discrete amount of time is drawn from a 8 million year time span and the lineages are allowed to evolve. At the end of the randomly chosen time period the divergence from the original population is measured. I then plotted these 5000 data points on both a log 10 scale and on the original scale of years.

## Acknowledgement

I am grateful to Erik Svensson for comments on earlier drafts of this manuscript.

